# Hierarchical gradients of multiple timescales in the mammalian forebrain

**DOI:** 10.1101/2023.05.12.540610

**Authors:** Min Song, Eun Ju Shin, Hyojung Seo, Alireza Soltani, Nicholas A. Steinmetz, Daeyeol Lee, Min Whan Jung, Se-Bum Paik

**Author notes:** M.S. and E.J.S. contributed equally to this work. **Author Contributions:** S.P., D.L., and M.W.J. designed and supervised the project. E.J.S. curated the previously published or publicly available data. M.S. analyzed the data. H.S. advised on the code for the autoregressive model. M.S. and E.J.S. wrote the first draft of the manuscript and created the figures. S.P., D.L., and M.W.J. edited the manuscript. All authors discussed and commented on the manuscript.

## Abstract

Many anatomical and physiological features of cortical circuits, ranging from the biophysical properties of synapses to the connectivity patterns among different neuron types, exhibit consistent variation along the hierarchical axis from sensory to association areas. Notably, the scale of temporal correlation of neural activity at rest, known as the intrinsic timescale, increases systematically along this hierarchy in both primates and rodents, analogous to the growing scale and complexity of spatial receptive fields. However, how the timescales for task-related activity vary across brain regions and whether their hierarchical organization appears consistently across different mammalian species remain unexplored. Here, we show that both the intrinsic timescale and the timescales of task-related activity follow a similar hierarchical gradient in the cortices of monkeys, rats, and mice. We also found that these timescales covary similarly in both the cortex and basal ganglia, whereas the timescales of thalamic activity are shorter than cortical timescales and do not conform to the hierarchical order predicted by their cortical projections. These results suggest that the hierarchical gradient of cortical timescales might be a universal feature of intra-cortical circuits in the mammalian brain.

**Significance Statement:** A gradual increase in the intrinsic timescales of cortical activity along the anatomical hierarchy reflects the functional specialization of cortical circuits. However, it is unknown whether this gradient of timescales is a common feature across distinct mammalian species in both intrinsic and task-related timescales and whether it is also observed in subcortical areas. This study reveals that the hierarchical gradient of multiple cortical timescales is conserved across multiple mammalian species. By contrast, thalamic timescales were shorter than cortical timescales and did not follow the hierarchical order inferred from their cortical projections. These findings imply a crucial role of intra-cortical connections in structuring distinct temporal dynamics observed across the cortex.

## Introduction

Compared to the peripheral sensory receptors, neurons in the cerebral cortex are tuned to more complex spatial and temporal features of the animal’s environment (1–3). In addition, the window for spatial and temporal integration systematically increases along the anatomical hierarchy of the cortex. Consequently, compared to the neurons in the primary sensory cortices, neurons in the association cortical areas tend to have larger receptive fields and their activity is affected by sensory stimuli over a longer temporal window (4–10). Many other anatomical and physiological properties of cortical circuits also display coordinated changes along this cortical hierarchy (11–14). Notably, both electrophysiological recordings and neuroimaging studies based on the blood-oxygen-level-dependent (BOLD) signals have identified a similar hierarchical gradient in the intrinsic timescale—the timescale of spontaneous or resting-state activity—across different cortical areas in humans, monkeys, and rodents (10, 13–22).

Although previous studies have mostly focused on the variation in intrinsic timescales, neurons in higher-order cortical areas, such as the prefrontal cortex, often modulate their activity according to multiple task variables (23). This raises the possibility that a single neuron may display multiple timescales for intrinsic and other task-related activities. Indeed, in the primate cortex, the timescales of task-related signals, such as the animal’s choice and reward, are decoupled from the intrinsic timescales of the same neurons within a given cortical area but still systematically increase along the cortical hierarchy (19, 24). In the present study, we tested whether cortical neurons in rats and mice also display a hierarchical gradient in multiple timescales, as previously shown in non-human primates, and whether the relationship among these distinct timescales generalizes to the subcortical structures such as the thalamus and basal ganglia. We found that the timescales of task-related activity as well as the intrinsic timescales follow the same cortical hierarchy in all three species. We also found that timescales of cortical activity did not vary across layers. In addition, the timescales of thalamic activity were shorter and did not follow the hierarchical pattern expected from their cortical projections. These findings suggest that the hierarchical organization of cortical timescales reflects a universal feature of intra-cortical connections in mammalian species.

## Results

### Multiple timescales in the cortex follow anatomical hierarchy

We estimated four distinct types of timescales using a previously described autoregressive time-series model [(19); see Materials and Methods for details]. Two of these timescales, intrinsic and seasonal timescales, quantify how quickly the strength of temporally correlated neural activity decays within a trial and across trials, respectively (Fig. 1*A*, top). The other two, choice- and reward-memory timescales, describe how rapidly neural signals related to an animal’s choice and reward decay (Fig. 1*A*, middle and bottom). We analyzed data from monkeys [866 neurons; (24–28)], rats [4,972 neurons; (29–34)], and mice [11,139 neurons; (35)] performing different behavioral tasks, namely a matching pennies task, dynamic foraging, and visual discrimination, respectively (Fig. 1 *B-D*; see Materials and Methods for details, and SI Appendix, Table S1 for information about the regions analyzed and corresponding abbreviations). In all three tasks, the animal’s choices and rewards varied sufficiently across trials so that the time course of neural signals related to those behavioral variables could be independently assessed. We then examined how these timescales — intrinsic, seasonal, choice-memory, and reward-memory — vary with the anatomical hierarchy, which was previously estimated in monkeys (36–38) and mice (39) using the ratio of feedforward and feedback connections [hierarchy score; (38, 39)]. Given that mice and rats belong to the same subfamily, *Murinae* (40), we used the mice’s hierarchy scores as a proxy for the rats’ scores.

**Figure 1.**
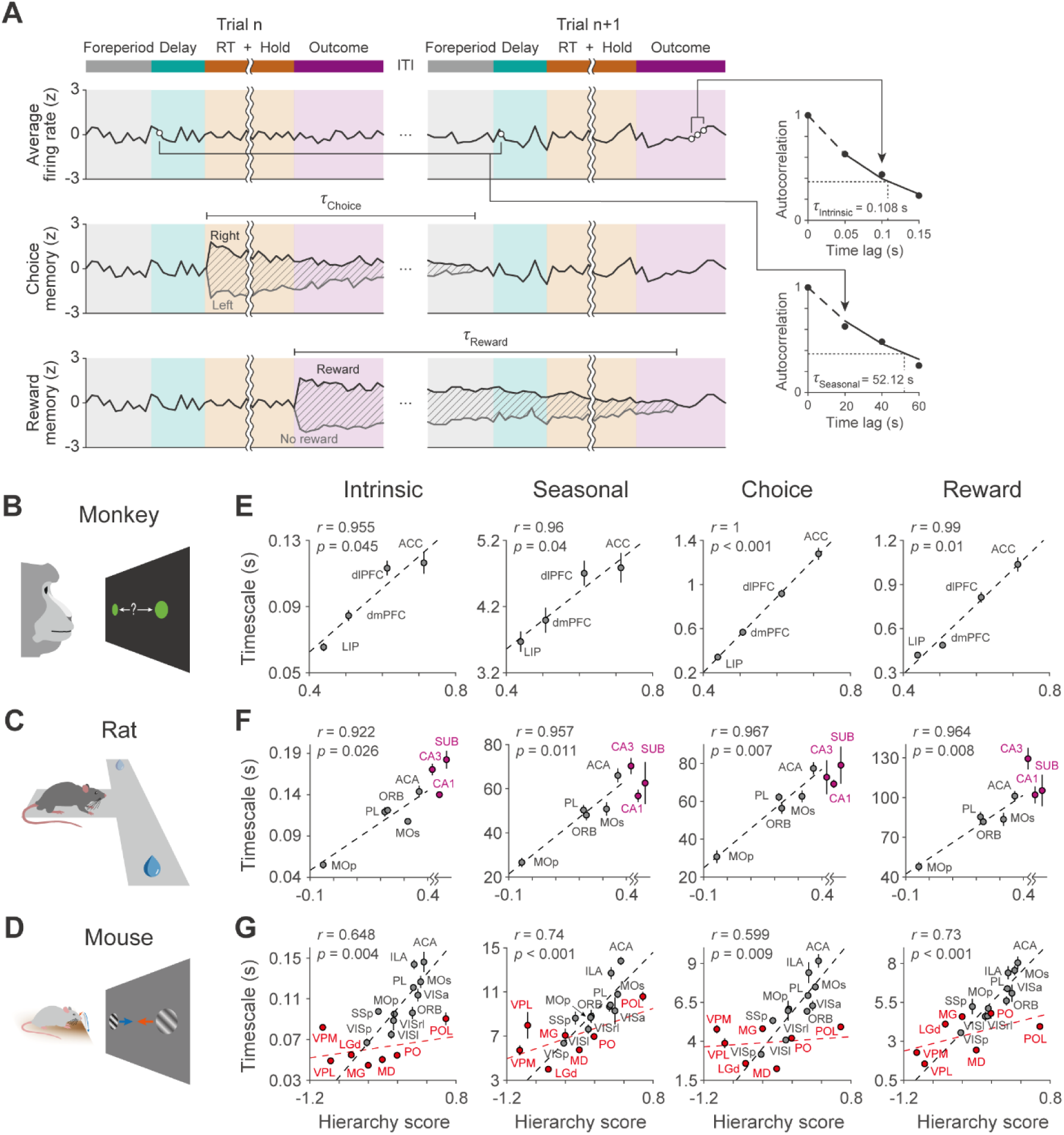
Multiple timescales in the neocortex follow the same hierarchical gradient. *(A)* Estimation of multiple timescales using an autoregressive model is illustrated with hypothetical neural data. Intrinsic and seasonal timescales estimate the rate of change in correlated neural activity within and across trials, respectively, whereas choice- and reward-memory timescales quantify the decay of neural signals related to the animal’s choice and its outcome, respectively. (*B*-*D*) Behavioral task for each species. Matching pennies task for monkeys (*B*), dynamic foraging task for rats (*C*), and visual discrimination task for mice (*D*). (*E*-*G*) Correlation between the timescales of neural activity and anatomical hierarchy score in monkeys (*E*), rats (*F*), and mice (*G*). Each dot and error bar respectively indicate the median and standard error of the timescale within each area. In (*F*), black and pink dots represent data from the neocortex and hippocampus, respectively. Note that the data from the hippocampus (CA3, CA1, and SUB) do not have a precise hierarchy score. In (*G*), black and red dots represent data from the neocortex and thalamus, respectively. Dashed lines denote linear regression. The Pearson correlation coefficients (r) and the p values are shown for the data pooled from all areas in each species.

We found that multiple timescales in cortical activity increased with the anatomical hierarchy score in all three species. As previously reported (19), intrinsic, seasonal, choice-memory, and reward-memory timescales all increased with the anatomical hierarchy score in the monkey cortex (Fig. 1*E*). Similarly, each of these timescales also increased significantly along the cortical hierarchy in both rats (Fig. 1*F*, black dots) and mice (Fig. 1*G*, black dots). This suggests that the hierarchical ordering of timescales is a common feature of the cortical organization across mammalian species. We also tested whether timescales of neural activity may vary across cortical layers (8, 41) in mice and found that none of the timescales analyzed in our study significantly varied across different layers and consistently followed the same anatomical hierarchy (Fig. 2 and SI Appendix, Table S2). This finding further underscores the contribution of intra-cortical circuits to the hierarchical organization of timescales in cortical activity.

**Figure 2.**
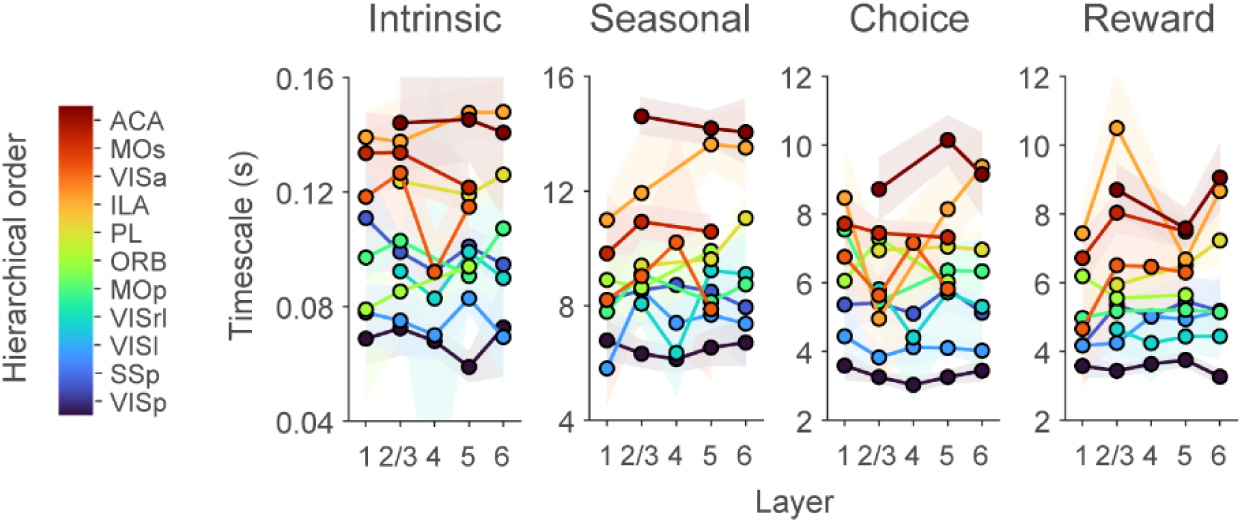
Cortical timescales are consistent across cortical layers. The intrinsic, seasonal, choice-memory, and reward-memory timescales are shown for each cortical layer in mice. To ensure the reliability of the analysis, only layers with more than 10 neurons and areas containing both superficial (layers 2/3 and 4) and deep layer (layers 5 and 6) neurons were analyzed. Note that layer 4 was not included in accordance with the Allen Brain Reference Atlas in the frontal cortex (MOp, MOs, ORB, PL, ILA, and ACA). Colored circles and shaded areas indicate the median and the standard error of timescales within each cortical layer, respectively.

Although the data from rats included neurons recorded in the CA1 and CA3 regions of the hippocampus and the subiculum, precise hierarchical scores for these regions are not available. However, it is reasonable to assume that the hippocampus and subiculum are at the top of the cortical hierarchy, as they receive neocortical inputs through the superficial layer of the entorhinal cortex and project to the deep layer of that in both monkeys and rodents (36, 42, 43). We found that the timescales of the hippocampus and subiculum were significantly longer than those of the neocortex (Fig. 1*F*, pink dots, Wilcoxon rank-sum test; |z| > 6.35 and p < 10^−9^ for all timescales). This suggests that the hierarchical gradients of multiple timescales in the cortex might extend to the hippocampal formation.

### Thalamic timescales are short and homogeneous

The hierarchical gradient of cortical timescales may stem from the local recurrent connections within each cortical area (44) or might rely on subcortical structures strongly connected to the cortex, such as the thalamus (45, 46). To distinguish between these possible scenarios, we analyzed the activity recorded from the mouse thalamus and tested whether timescales in various thalamic nuclei mirror the hierarchical gradient of their cortical counterparts (39). For each of the four timescales analyzed, we found that the median timescale in the thalamus was significantly shorter than that in the cortex (Fig. 1*G*, red dots, Wilcoxon rank-sum test; |z| > 2.35, p < 0.05, for all timescales) and was not significantly correlated with the hierarchy scores of their cortical counterparts (Pearson correlation coefficient, Intrinsic, *r* = 0.343, p = 0.452; Seasonal, r = 0.635, p = 0.126; Choice, r = 0.172; p = 0.713; Reward, r = 0.55, p = 0.201; see SI Appendix, Table S3 for information about differences in the correlations between the cortex and thalamus). These results suggest that thalamic afferents may not be a major contributor to the hierarchical gradient of cortical timescales.

### Multiple timescales covary globally, but not locally, throughout the brain

We next examined whether various task-related timescales correlate with intrinsic timescales. As previously shown in monkeys (Fig. 3*A*), seasonal, choice-memory, and reward-memory timescales were all significantly correlated with intrinsic timescales across different neocortical areas in rats (Fig. 3*B*, black dashed line; Pearson correlation coefficient, *r* > 0.95 and p < 0.01 for all timescales) and mice (Fig. 3*C*, black dashed line; Pearson correlation coefficient, *r* > 0.95 and p < 10^−5^ for all timescales). All these correlations were still statistically significant when the timescales in the rat hippocampus and striatum and the mouse thalamus were also included (Fig. 3B and 3C). We also analyzed the data from the striatum and hippocampus in rats (Fig. 3*B*, green and pink dots) and the thalamus in mice (Fig. 3*C*, red dots) separately, and found significant correlations in rats (Fig. 3*B*, orange dashed line; *r* > 0.93 and p < 0.01 for all timescales; SI Appendix, Table S4, Region×Intrinsic), but not in mice (Fig. 3*C*, orange dashed line; Pearson correlation coefficient, Seasonal, *r* = 0.44, p = 0.32; Choice, *r* = 0.49, p = 0.263; Reward, r = −0.056, p = 0.91; SI Appendix, Table S4, Region×Intrinsic). The lack of correlation among multiple timescales in the thalamus might be due to the lack of sufficient variability across different nuclei.

**Figure 3.**
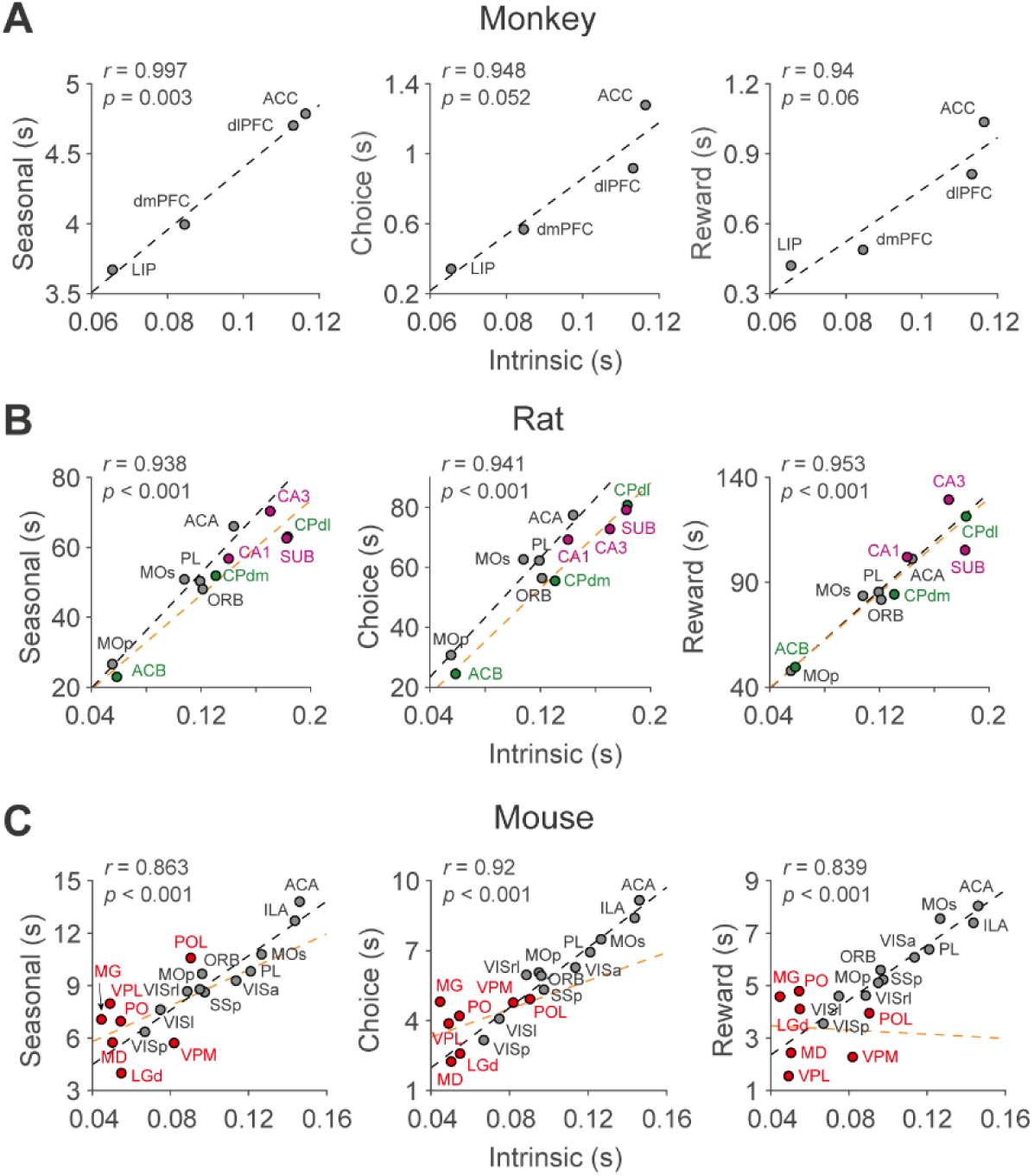
Multiple timescales are correlated across brain areas. (*A*-*C*) Correlation between the intrinsic timescale and the seasonal, choice-memory, and reward-memory timescales in monkeys (*A*), rats (*B*), and mice (*C*). Median timescales are shown for the neocortex (black), striatum (green), hippocampus (pink), and thalamus (red). Dashed lines show the regression outcomes for the neocortical (black) and non-neocortical (orange) areas. The Pearson correlation coefficients (r) and the p values for the data pooled from all areas are shown in each panel.

The above results suggest that multiple timescales in the mammalian cortex might be governed by a single mechanism, such as the systematic variation in the strength of recurrent connectivity across different brain areas (44, 47), or by multiple mechanisms that covary (48, 49). If so, different timescales of individual neurons would be correlated with one another, even within a given brain region. However, we found that different timescales of individual neurons were not significantly correlated with one another within any brain area in rats or mice (p > 0.05, after Bonferroni correction for multiple comparisons; SI Appendix, Fig. S1*A*), consistent with a previous finding in the primate cortex (19). To examine this further, we pooled the timescales of individual neurons from all brain areas in each species after subtracting the median of the timescale in each area. We found that different timescales were still not significantly correlated in any species, regardless of whether the results from the neocortex, striatum, hippocampus, and thalamus were analyzed together or separately (p > 0.05, after Bonferroni correction for multiple comparisons; SI Appendix, Fig. S1*B*). These results suggest that the hierarchical gradient of multiple timescales is shaped by multiple factors that vary consistently across brain areas but independently within the population of neurons in each brain area.

## Discussion

Previous studies have demonstrated a hierarchical gradient of intrinsic timescales in resting or spontaneous cortical activity in humans (13, 14, 20), monkeys (15–19, 21, 22), and rodents (10) using electrophysiological recordings and neuroimaging data. In the present study, we characterized the timescales of task-relevant and task-irrelevant spiking activity across diverse brain areas in monkeys, rats, and mice using the same analysis framework. Our analyses revealed that the parallel, independent progression of multiple timescales along the cortical hierarchy is preserved in all three species and, at least in mice, did not vary across different cortical layers. Although precise hierarchical scores were not available for the rat hippocampus and basal ganglia, we found that the multiple timescales in these brain areas follow a relationship similar to that observed among multiple cortical timescales.

### Hierarchical organization of timescales in cortex

Most studies on the timescales of cortical activity have focused on the intrinsic timescale, which refers to the rate at which the autocorrelation in spontaneous neural activity decays over time (15, 16). Intrinsic timescales measured from spiking activity, field potentials, and BOLD signals consistently show a gradual increase from primary sensory cortical areas to higher-order association cortical areas, such as the anterior cingulate cortex (13, 14, 16, 20–22). This pattern is often interpreted as reflecting the different computational needs of sensory versus association cortical regions (6, 7). Neurons in sensory cortical areas need to process incoming signals rapidly to represent dynamics in the sensory environment with high temporal resolution. In contrast, neurons in higher-order cortical areas may specialize in extracting more stable features of the environment.

Estimates of intrinsic timescales might depend on the nature of the neural signals analyzed. Indeed, the values of intrinsic timescales reported in previous studies vary substantially with the type of neural signals, ranging from tens of milliseconds in electroencephalography (EEG) and electrocorticography (ECoG) recordings (14, 22), to hundreds of milliseconds in spiking activity (10, 15–19) and up to several seconds in BOLD signals (13, 20, 21). Differences between intrinsic timescales estimated from EEG/ECoG and spiking activity might be reconciled by computational studies showing that small differences in the dynamic properties of the excitatory synapses underlying local recurrent connections across different regions can lead to large variations in the intrinsic timescales reflected in spiking activity (47, 50, 51). Similarly, the larger estimates of intrinsic timescales in fMRI studies might be partly due to the filtering properties of the hemodynamic response that links underlying neural activity with BOLD signals (52).

Our results also suggest that it is crucial to consider the presence of multiple timescales, including those specifically related to tasks and animal’s behavior, to fully understand the relationship between timescales estimated under different conditions across studies. For example, we observed that timescales spanning multiple trials (seasonal timescale) and timescales associated with the animal’s choice and reward (choice- and reward-memory timescales) are much longer than intrinsic timescales. The consistency of these relationships across three different species indicates their robustness, despite variations in the specifics of behavioral tasks. However, choice- and reward-memory timescales may vary flexibly with task demands and the animal’s behavioral states (19), which could potentially confound estimates of intrinsic timescales if different timescales are not properly accounted for with appropriate models. Similarly, the relatively long intrinsic timescales reported in fMRI studies might reflect long-range temporal correlations more akin to seasonal timescales than the intrinsic timescale observed in spiking activity.

### Timescales beyond the cortex

Although previous studies on the intrinsic timescales of neural activity have mostly focused on the cortex, some have extended these analyses to subcortical structures and the hippocampus. For example, the intrinsic timescale of activity in first-order sensory nuclei of the thalamus, such as the lateral geniculate nucleus (LGN) and ventral posterior lateral nucleus (VPL), is similar to those observed in the primary sensory cortical areas (10, 53). However, since these studies did not examine timescales for higher-order thalamic nuclei, it was unclear whether the timescales of neural activity in the thalamus follow a hierarchical gradient similar to their cortical counterparts.

Our results indicate that thalamic timescales are consistently shorter and do not exhibit a hierarchical pattern. In addition, the observation that stimulus-driven or persistent activity in the cortex can be rapidly interrupted by inhibiting thalamic inputs (54, 55) suggests that the contribution of thalamic inputs to driving cortical activity may be multiplicative, functioning to gate rather than generate cortical signals with specific timescales (56–58). These results imply that the sources of hierarchical gradients in the cortex may reside within the cortex itself.

The hippocampus is commonly assumed to be at the top of the anatomical hierarchy (36, 42, 43), although its precise hierarchy score is not available for quantitative analyses in the rats. A previous study of human EEG recordings reported that intrinsic timescales increased from the parietal cortex to the prefrontal cortex, reaching their maximum in the medial temporal lobe (14). Similarly, we found that multiple timescales in the hippocampus and subiculum are longer than those in other neocortical areas, suggesting that the hierarchical gradient of cortical timescales extends to the hippocampal formation.

Similar to the research on thalamic and hippocampal timescales, only a few studies have reported intrinsic timescales in the basal ganglia. Some fMRI studies have suggested that timescales in the striatum may reflect those of the connected cortical areas (20, 21). However, the anatomical distribution of timescales reported in these studies has been inconsistent. For instance, Raut et al. (2020) found longer timescales in the ventral striatum compared to the putamen, whereas Manea et al. (2022) reported the shortest timescales in the ventral striatum. In our study, all types of timescales were consistently shorter in the ventral striatum than in the dorsal striatum, which aligns with the findings of Manea et al. (2022). However, significant differences in the anatomical organization of the striatum between primates and rodents may exist (59, 60). Additionally, differences between cortical and subcortical hemodynamic responses could influence the relationship between timescales estimated from neural activity and those derived from BOLD signals. Therefore, direct comparisons of subcortical timescales across studies should be approached with caution.

### Factors contributing to the gradient of timescales

The observation that multiple timescales identified in both intrinsic and task-related neural activity increase similarly along the anatomical hierarchy in the cortex across different species suggests a universal organizational principle. Previous studies have identified similar gradients in various network, cellular, and molecular properties along the sensory-to-association or unimodal-to-transmodal axis of the neocortex (11–14, 48, 49). Understanding how these factors interact to produce consistent gradients in multiple timescales without correlation within each cortical area might be challenging. While previous modeling works have provided insights into this issue, they have typically focused on a single gradient of intrinsic timescales along the cortical hierarchy.

Wang and his colleagues have suggested that neuronal timescales are primarily determined by local connectivity, particularly the dense recurrent connections in cortical circuits (47, 51, 61, 62). Their work demonstrates how synaptic properties and connectivity patterns can shape neural timescales by modulating the balance between excitatory and inhibitory inputs. On the other hand, other modeling studies suggest that variations in timescales may arise from multiple factors, whose contributions can differ across brain areas. For example, neural circuits with heterogeneous assemblies can generate timescales that span several orders of magnitude, depending on assembly size (63). This offers a plausible explanation for the wide variation in timescales observed in cortical activity, which can be decorrelated within each cortical area. These model results suggest that multiple factors contribute to the organization and modulation of neuronal timescales, emphasizing the need for considering these factors collectively in future models.

In conclusion, our findings highlight the hierarchical gradients of multiple timescales as a fundamental aspect of cortical dynamics. We propose that intra-cortical connections primarily modulate cortical timescales, while additional mechanisms may influence subcortical timescales. Further investigations into multiple timescales across different brain regions and species could elucidate the complex relationship between brain connectivity and neuronal temporal dynamics.

### Materials and Methods The Neural data

We analyzed single-neural activity collected previously from multiple brain regions of monkeys, rats, and mice. Specifically, we analyzed activity from four cortical areas in monkeys (24–28) and from five cortical areas, three striatal subregions, and three hippocampal regions in rats (29–34).

In addition, we used the dataset for mice publicly available at https://figshare.com/articles/dataset/Dataset_from_Steinmetz_et_al_2019 /9598406 (35), which includes neural activity recorded from eleven cortical areas and seven thalamic nuclei. The numbers of neurons in each area from the three species are listed in SI Appendix, Table S1.

### Behavioral task

Monkeys performed a matching pennies task (24–28). The trial started with a yellow square presented at the center of the computer screen. After a delay of 0.5 seconds, two green disks were presented horizontally. The animal shifted its gaze towards one of the targets within one second and maintained its fixation for 0.5 seconds. Subsequently, a red ring appeared around the target selected by the computer opponent, and the animal was rewarded only when it chose the same target as the computer. The computer opponent was programmed to predict the animal’s choice based on the animal’s previous choices and reward outcomes.

Rats performed dynamic foraging tasks (29–34). The rat was presented with two goal locations and allowed freely to choose one of them to receive a reward. One of the four reward probability pairs for left and right goals (left:right = 0.72:0.12, 0.63:0.21, 0.12:0.72, or 0.21:0.63) was used in each block with the number of trials in each block randomly determined.

Mice performed a visual discrimination task (35). The trial began with visual stimuli presented on the left and right sides of the computer screen. After a delay of 0.5 to 1.2 seconds followed by an auditory cue, the animal moved a stimulus located at either side to the center of the screen by turning the wheel. If the animal chose the stimulus with higher contrast, it was rewarded. If the two stimuli were of equal contrast, the animal was rewarded with 50% probability for a left or right choice. If no stimuli were presented on the screen, the animal was rewarded only when it did not turn the wheel for 1.5 seconds following the auditory tone cue.

### Hierarchy scores

The anatomical hierarchy scores for monkeys and mice used in this study were obtained from two prior studies (38, 39), and they are publicly available at https://balsa.wustl.edu/ and https://github.com/AllenInstitute/MouseBrainHierarchy, respectively. For the mice data, we used the hierarchy score calculated from the feedforward and feedback connections across the neocortical and thalamic areas, which are indicated as the ‘CC+TC+CT iterate scores in the result file ‘hiearchy_summary_CreConf.xlsx’, to compare the timescales from the areas in the neocortex and thalamic nuclei. For the rat data, the hierarchy scores from the mice were used.

### Classification of cortical layers in mice

Since the neural data from mice were recorded with Neuropixels probes (64), we mapped cortical neurons to each cortical layer according to their channel position detected at peak amplitude in the Allen Mouse Brain Common Coordinate Framework (65).

### Autoregressive model

Neural activity recorded from animals during the behavior tasks displays ongoing fluctuations within and across trials in addition to changes related to the animal’s choices and their outcomes. In the present study, the timescales of intrinsic changes in the neural activity within a single trial and across multiple trials, referred to as ‘intrinsic’ and ‘seasonal’ timescales, respectively, were estimated using two separate groups of terms in a third-order autoregressive (AR) model (19). In addition, we estimated the timescales of neural signals related to the animal’s choice and its outcome by modeling them with the sum of the exponential functions with different time constants (Fig. 1*A*). To calculate these two timescales simultaneously, we used the same autoregressive model as employed in a previous study (19), where the timescales were estimated with a fitting algorithm to predict the current spike count of a neuron using the previous neural response and the history of the animal’s choices and their outcomes in three previous trials. In this model, the spike counts in each time bin are predicted using the following model:

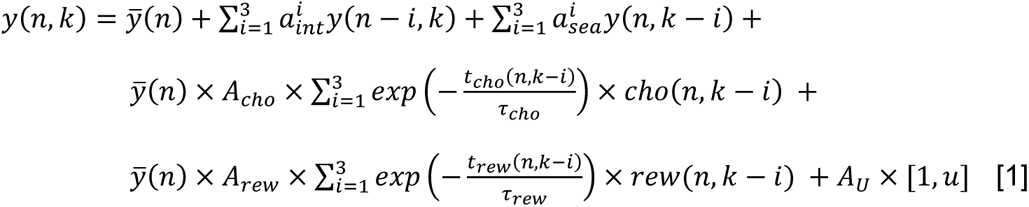

where y(n, k) is the spike count in the n^th^ bin (with a resolution of 50ms) of the k^th^ trial and *ȳ*(𝑛) is a constant term for each bin and 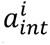 and 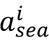 are the corresponding coefficients for the intrinsic and seasonal AR components. These results represent the intrinsic and seasonal fluctuations in the neural activity. 𝑡_𝑐ℎ𝑜_(𝑛, 𝑘) and 𝑡*_rew_*(𝑛, 𝑘) denote the time difference between the current time and the time when the choice was made and the reward was given in prior trials, respectively (with a resolution of 50ms). The free parameters of the model are the autoregressive coefficients for the intrinsic 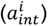 and seasonal 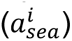 timescales; the amplitudes of the choice-memory and reward-memory components are denoted respectively by 𝐴_𝑐ℎ𝑜_ and 𝐴*_rew_*; and the timescales of the choice and reward signals are likewise indicated by 𝜏_𝑐ℎ𝑜_ and 𝜏*_rew_*. 𝑢 and 𝐴_𝑈_ represent the vectors of behavioral terms (current choice, reward, and their interaction (choice×reward)) and their coefficients, respectively. The construction of the regressor vector, 𝑢, is tailored to the specific behavioral paradigms employed. For monkeys, u is composed of three choice-related regressors capturing the intervals [0, 500] milliseconds post-onset of choice targets, target fixation, and reward feedback, along with a single reward regressor and an interaction regressor delineating the [0, 500] milliseconds following reward feedback. In mice, u was defined similarly, including three choice-related regressors for the periods following the onset of a visual stimulus, an auditory tone cue, and reward feedback, complemented by one reward regressor and an interaction regressor for the interval immediately after reward feedback. For rats, the regressor vector included two choice-related regressors, covering the time [0, 500] milliseconds after initiating an approach towards the reward and the receipt of reward feedback, in addition to a single reward and an interaction regressor for the post-reward feedback phase. We estimated the time constants of choice- and reward-memory effects from the corresponding exponential functions.

The timescale of the intrinsic and seasonal fluctuations was calculated from the eigenvalues of three AR coefficients, as follows:

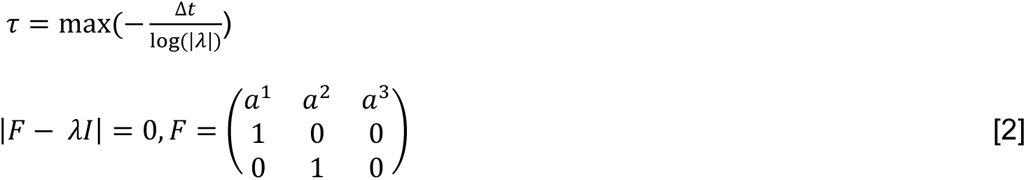

where Δt is the size of the time bin for each component (Δ𝑡 = 50ms for the intrinsic timescale and the average trial length for the seasonal timescale).

To assess the statistical significance of each timescale, we compared log likelihood values from the full ARX model (Eq. **1**) and from the model without a term that corresponds to the timescale in question. The difference in log likelihood values between these two models provided a measure of the contribution of the tested timescale on the ARX model. The statistical significance of this difference was tested using a Chi-square test. We found that distributions of timescales from all neurons were not significantly different from those only including the neurons with statistically significant terms (Kolmogorov-Smirnov test, p > 0.05, after Bonferroni correction for multiple comparisons), and therefore all neurons with mean firing rate > 0.1 spikes/s are included in the analyses.

### Multiple-regression models

To examine the difference between the hierarchical gradient of each timescale in superficial and deep layers, we used the following regression model (SI Appendix, Table S2):

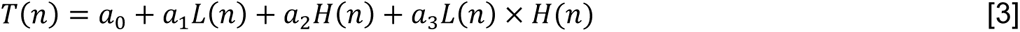

where 𝑇(𝑛) is the median of timescale in the superficial or deep layer of n^th^ brain area, 𝐿(𝑛) is a dummy variable to indicate whether each layer belongs to the superficial or deep layer (that is, 0 and 1 for the superficial and deep layer, respectively), 𝐻(𝑛) is the hierarchy score, and 𝐿(𝑛) × 𝐻(𝑛) is the interaction term. Thus, the null hypothesis that 𝑎_3_ = 0 implies that the strength of the correlation between the timescale and the hierarchy score is not different for the two layers.

Similarly, to examine the difference between the hierarchical gradient of each timescale in the cortex and thalamus, we used the following regression model (SI Appendix, Table S3):

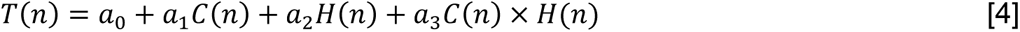

where 𝑇(𝑛) is the median of timescale in the n^th^ brain area, 𝐶(𝑛) is a dummy variable to indicate whether or not each area belongs to the cortex (that is, 0 and 1 for the cortex and thalamus, respectively).

Additionally, to test whether the correlation among multiple types of timescales differs for the neocortical and non-neocortical areas, we used the following regression model (SI Appendix, Table S4):

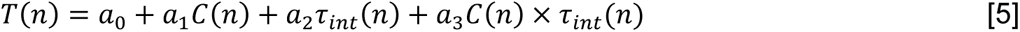

where 𝑇(𝑛) is the median of timescale in the n^th^ brain area, 𝐶(𝑛) is a dummy variable to indicate whether or not each area belongs to the cortex (that is, 0 and 1 for the neocortical and non-neocortical area, respectively).

### Quantification and statistical analysis

Pearson’s correlation coefficient was used to determine the correlation between the hierarchical score and multiple timescales (Fig. 1*E*-*G*), and between the intrinsic timescale and other timescales (Fig. 3 and SI Appendix, Fig. S1). The Wilcoxon rank-sum test (Mann-Whitney U test) was used to determine the difference in the timescale values between the cortical and hippocampus in rats and the thalamic areas in mice.

## Acknowledgments

This work was supported by a grant from the National Research Foundation of Korea funded by the Korean government NRF-2022R1A2C3008991 (S.P.), the Singularity Professor Research Project of KAIST (S.P.), the National Institute of Health MH137210 (D.L), DA047870 (A.S.), and the Research Center Program of the Institute for Basic Science IBS-R002-A1 (M.W.J.).

---

**Fig. S1.**
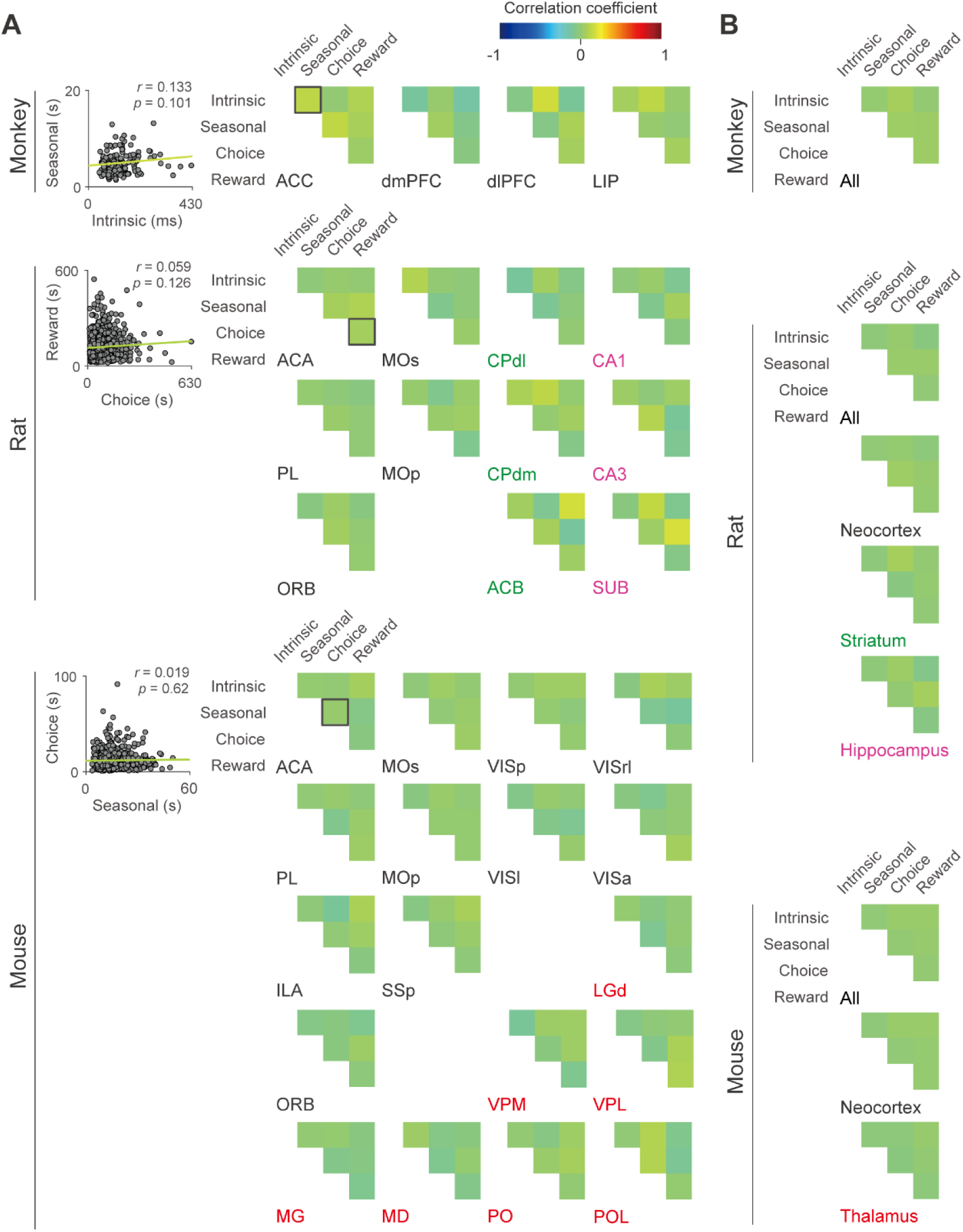
Lack of a correlation between multiple timescales across individual neurons. (A) Pearson correlation coefficients between all pairs of different timescales measured from individual neurons in different brain areas of monkeys, rats, and mice. For each species, an example of the relationship between different timescales across neurons is shown for the anterior cingulate cortex (ACC or ACA), as indicated by the black box in the correlation matrix. Solid lines represent linear regressions of the data. (B) Corresponding results for the residual timescales combined for all brain areas tested in the present study. None of these correlations were statistically significant (p > 0.05, after Bonferroni correction for multiple comparisons).

**Table S1.** Brain regions analyzed in each species.

| Species | Abbreviation | Full name | Number of neurons | Number of animals |
| --- | --- | --- | --- | --- |
| Monkey | ACC | Anterior cingulate cortex | 154 | 2 |
|  | dIPFC | Dorsolateral prefrontal cortex | 189 | 5 |
|  | dmPFC | Dorsomedial prefrontal cortex | 185 | 2 |
|  | LIP | Lateral intraparietal cortex | 205 | 3 |
|  | Total |  | 866 | 6 |
| Rat | ACA | Anterior cingulate area | 679 | 5 |
|  | ORB | Orbital area | 1148 | 3 |
|  | PL | Prelimbic area | 854 | 6 |
|  | MOp | Primary motor area | 227 | 3 |
|  | MOs | Secondary motor area | 411 | 3 |
|  | CPdm | Dorsomedial caudoputamen | 414 | 6 |
|  | CPdl | Dorsolateral caudoputamen | 129 | 3 |
|  | ACB | Nucleus accumbens | 165 | 3 |
|  | CA1 | Field CA1 (Cornu Ammonis 1) | 599 | 9 |
|  | CA3 | Field CA3 (Cornu Ammonis 3) | 231 | 4 |
|  | SUB | Subiculum | 115 | 3 |
|  | Total |  | 4972 | 27 |
| Mouse | ACA | Anterior cingulate area | 841 | 8 |
|  | ORB | Orbital area | 660 | 4 |
|  | PL | Prelimbic area | 863 | 7 |
|  | ILA | Infralimbic area | 411 | 3 |
|  | MOp | Primary motor area | 793 | 3 |
|  | MOs | Secondary motor area | 1743 | 9 |
|  | SSp | Primary somatosensory area | 515 | 4 |
|  | VISa | Anterior visual area | 340 | 4 |
|  | VISI | Lateral visual area | 434 | 2 |
|  | VISp | Primary visual area | 1118 | 8 |
|  | VISrl | Rostrolateral visual area | 286 | 2 |
|  | LGd | Dorsal part of the lateral geniculate complex | 882 | 5 |
|  | POL | Posterior limiting nucleus of the thalamus | 215 | 3 |
|  | MD | Mediodorsal nucleus of the thalamus | 397 | 3 |
|  | VPL | Ventral posterolateral nucleus of the thalamus | 301 | 3 |
|  | PO | Posterior complex of the thalamus | 692 | 4 |
|  | VPM | Ventral posteromedial nucleus of the thalamus | 353 | 2 |
|  | MG | Medial geniculate complex of the thalamus | 295 | 2 |
|  | Total |  | 11139 | 10 |

The number of neurons and animals obtained from each region and species used for analysis are shown.

**Table S2.** Results from the regression model to test the differences in the correlations between the timescales for the cortical layers in mice.

| **Timescales** | **Intercept** | **Layer** | **Hierarchy score** | **Layer×Hierarchy score** |
| --- | --- | --- | --- | --- |
| **Intrinsic** | 20.054 ($9.18 \times 10^{-13}$) | −0.078 (0.939) | 3.703 (0.002) | 0.171 (0.867) |
| **Seasonal** | 17.884 ($5.33 \times 10^{-12}$) | −0.126 (0.902) | 3.227 (0.005) | −0.15 (0.882) |
| **Choice** | 18.413 ($3.41 \times 10^{-12}$) | 0.214 (0.834) | 4.839 ($1.81 \times 10^{-4}$) | −0.265 (0.795) |
| **Reward** | 21.796 ($2.53 \times 10^{-13}$) | 0.015 (0.988) | 5.274 ($7.57 \times 10^{-5}$) | −0.519 (0.611) |

Results of t-statistics (p) are shown for different regressors in the regression model given by Eq. 3 (see Materials and Methods for details). To increase the reliability of the statistical test, layers were grouped into superficial (layers 2/3 and 4) and deep (layers 5 and 6) layers.

**Table S3.**
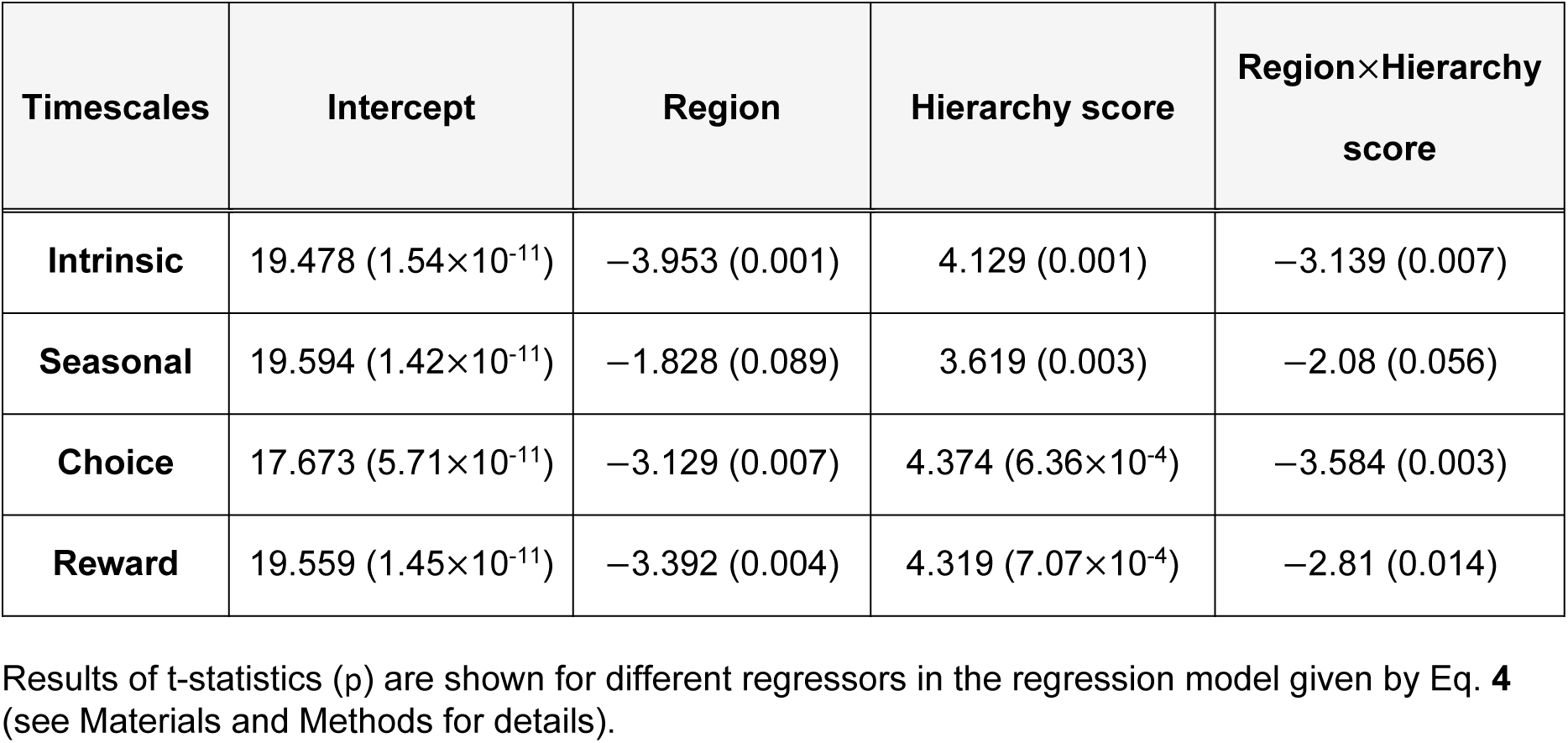
Results from the regression model to test the differences in the correlations between the timescales for the neocortex and thalamic nuclei in mice.

| **Timescales** | **Intercept** | **Region** | **Hierarchy score** | **Region×Hierarchy score** |
| --- | --- | --- | --- | --- |
| **Intrinsic** | 19.478 ($1.54 \times 10^{-11}$) | −3.953 (0.001) | 4.129 (0.001) | −3.139 (0.007) |
| **Seasonal** | 19.594 ($1.42 \times 10^{-11}$) | −1.828 (0.089) | 3.619 (0.003) | −2.08 (0.056) |
| **Choice** | 17.673 ($5.71 \times 10^{-11}$) | −3.129 (0.007) | 4.374 ($6.36 \times 10^{-4}$) | −3.584 (0.003) |
| **Reward** | 19.559 ($1.45 \times 10^{-11}$) | −3.392 (0.004) | 4.319 ($7.07 \times 10^{-4}$) | −2.81 (0.014) |

Results of t-statistics (p) are shown for different regressors in the regression model given by Eq. 4 (see Materials and Methods for details).

**Table S4.** Results from the regression model used to test the differences in the relationships between the intrinsic and other timescales between the neocortical and non-neocortical areas.

| Animal | Timescales | Intercept | Region | Intrinsic | Region×Intrinsic |
| --- | --- | --- | --- | --- | --- |
| Rat | Seasonal | 0.376 (0.718) | 0.241 (0.817) | 5.518 ($8.9 \times 10^{-4}$) | −0.847 (0.425) |
| | Choice | 0.428 (0.682) | −0.329 (0.752) | 6.832 ($2.46 \times 10^{-4}$) | −0.646 (0.539) |
|  | Reward | 1.008 (0.347) | 0.053 (0.959) | 4.146 (0.004) | −0.11 (0.916) |
| Mouse | Seasonal | 0.77 (0.454) | 0.904 (0.381) | 4.796 ($2.85 \times 10^{-4}$) | −0.763 (0.458) |
| | Choice | −0.628 (0.54) | 1.872 (0.082) | 7.283 ($4.02 \times 10^{-6}$) | −1.804 (0.093) |
| | Reward | 0.232 (0.82) | 1.955 (0.071) | 4.924 ($2.24 \times 10^{-4}$) | −2.474 (0.027) |

Results of t-statistics (p) are shown for different regressors in the regression model given by Eq. 5 (see Materials and Methods for details). Non-neocortical areas refer to the striatum and hippocampus for rats, and the thalamus for mice.

## References

1. D. H. Hubel, T. N. Wiesel, Receptive fields, binocular interaction and functional architecture in the cat’s visual cortex. J. Physiol. 160, 106–154 (1962).

2. V. B. Mountcastle, W. H. Talbot, H. Sakata, J. Hyvärinen, Cortical neuronal mechanisms in flutter-vibration studied in unanesthetized monkeys. Neuronal periodicity and frequency discrimination. J. Neurophysiol. 32, 452–484 (1969).

3. N. Ulanovsky, L. Las, D. Farkas, I. Nelken, Multiple time scales of adaptation in auditory cortex neurons. J. Neurosci. 24, 10440–10453 (2004).

4. P. Lennie, Single units and visual cortical organization. Perception 27, 889–935 (1998).

5. U. Hasson et al., A hierarchy of temporal receptive windows in human cortex. J. Neurosci. 28, 2539–2550 (2008).

6. S. J. Kiebel, J. Daunizeau, K. J. Friston, A hierarchy of time-scales and the brain. PLoS Comput. Biol. 4, e1000209 (2008).

7. C. J. Honey et al., Slow cortical dynamics and the accumulation of information over long timescales. Neuron 76, 423–434 (2012).

8. S. E. Cavanagh, L. T. Hunt, S. W. Kennerley. A Diversity of Intrinsic Timescales Underlie Neural Computations. Front Neural Circuits. 14, 615626 (2020).

9. A. Soltani, J. D. Murray, H. Seo, D. Lee, Timescales of Cognition in the Brain. Curr. Opin. Behav. Sci. 41, 30–37 (2021).

10. J. H. Siegle et al., Survey of spiking in the mouse visual system reveals functional hierarchy. Nature 592, 86–92, (2021).

11. J. B. Burt et al., Hierarchy of transcriptomic specialization across human cortex captured by structural neuroimaging topography. Nat. Neurosci. 21, 1251–1259 (2018).

12. B. D. Fulcher, J. D. Murray, V. Zerbi, X. J. Wang, Multimodal gradients across mouse cortex. Proc. Natl. Acad. Sci. U.S.A. 116, 4689–4695 (2019).

13. T. Ito, L. J. Hearne, M. W. Cole, A cortical hierarchy of localized and distributed processes revealed via dissociation of task activations, connectivity changes, and intrinsic timescales. Neuroimage 221, 117141 (2020).

14. R. Gao, R. L. van den Brink, T. Pfeffer, B. Voytek, Neuronal timescales are functionally dynamic and shaped by cortical microarchitecture. Elife 9, e61277 (2020).

15. T. Ogawa, H. Komatsu, Differential temporal storage capacity in the baseline activity of neurons in macaque frontal eye field and area V4. J Neurophysiol. 103, 2433–2445 (2010).

16. J. D. Murray et al., A hierarchy of intrinsic timescales across primate cortex. Nat. Neurosci. 17, 1661–1663 (2014).

17. V. Fascianelli, S. Tsujimoto, E. Marcos, A. Genovesio, Autocorrelation Structure in the Macaque Dorsolateral, But not Orbital or Polar, Prefrontal Cortex Predicts Response-Coding Strength in a Visually Cued Strategy Task. Cerebral Cortex 29, 230–241 (2017).

18. D. F. Wasmuht et al., Intrinsic neuronal dynamics predict distinct functional roles during working memory. Nat. Commun. 9, 3499 (2018).

19. M. Spitmaan, H. Seo, D. Lee, A. Soltani, Multiple timescales of neural dynamics and integration of task-relevant signals across cortex. Proc. Natl. Acad. Sci. U.S.A. 117, 22522–22531 (2020).

20. R. V. Raut, A. Z. Snyder, M. E. Raichle, Hierarchical dynamics as a macroscopic organizing principle of the human brain. Proc Natl Acad Sci U S A. 117, 20890–20897 (2020).

21. A. M. G. Manea et al., Intrinsic timescales as an organizational principle of neural processing across the whole rhesus macaque brain. Elife 11, e75540 (2022).

22. A. M. G. Manea et al., Neural timescales reflect behavioral demands in freely moving rhesus macaques. Nat. Commun. 15, 2151 (2024).

23. K. M. Tye et al., Mixed selectivity: Cellular computations for complexity. Neuron 112, 2289–2303 (2024).

24. E. Trepka et al., Training-dependent gradients of timescales of neural dynamics in the primate prefrontal cortex and their contributions to working memory. J. Neurosci. 44, e2442212023 (2023).

25. D. J. Barraclough, M. L. Conroy, D. Lee, Prefrontal cortex and decision making in a mixed-strategy game. Nat. Neurosci. 7, 404–410 (2004).

26. H. Seo, D. Lee, Temporal filtering of reward signals in the dorsal anterior cingulate cortex during a mixed-strategy game. J. Neurosci. 27, 8366–8377 (2007).

27. H. Seo, D. J. Barraclough, D. Lee, Lateral intraparietal cortex and reinforcement learning during a mixed-strategy game. J. Neurosci. 29, 7278–7289 (2009).

28. C. H. Donahue, H. Seo, D. Lee, Cortical signals for rewarded actions and strategic exploration. Neuron 80, 223–234 (2013).

29. H. Kim et al., Role of striatum in updating values of chosen actions. J. Neurosci. 29, 14701–14712 (2009).

30. H. Kim, D. Lee, M. W. Jung, Signals for previous goal choice persist in the dorsomedial, but not dorsolateral striatum of rats. J. Neurosci. 33, 52–63 (2013).

31. J. H. Sul et al., Distinct roles of rodent orbitofrontal and medial prefrontal cortex in decision making. Neuron 66, 449–460 (2010).

32. J. H. Sul, S. Jo, D. Lee, M. W. Jung, Role of rodent secondary motor cortex in value-based action selection. Nat. Neurosci. 14, 1202–1208 (2011).

33. H. Lee et al., Hippocampal neural correlates for values of experienced events. J. Neurosci. 32, 15053–15065 (2012).

34. S. H. Lee et al., Neural Signals Related to Outcome Evaluation Are Stronger in CA1 than CA3. Front. Neural Circuits 7, 11–40 (2017).

35. N. A. Steinmetz, P. Zatka-Haas, M. Carandini, K. D. Harris, Distributed coding of choice, action and engagement across the mouse brain. Nature 576, 266–273 (2019).

36. D. J. Felleman, D. C. Van Essenm, Distributed hierarchical processing in the primate cerebral cortex. Cereb. Cortex 1, 1–47 (1991).

37. N. T. Markov et al., A weighted and directed interareal connectivity matrix for macaque cerebral cortex. Cereb. Cortex 24, 17–36 (2014).

38. J. B. Burt et al., Hierarchy of transcriptomic specialization across human cortex captured by structural neuroimaging topography. Nat. Neurosci. 21, 1251–1259 (2018).

39. J. A. Harris et al., Hierarchical organization of cortical and thalamic connectivity. Nature 575, 195–202 (2019).

40. P. H. Fabre, L. Hautier, D. Dimitrov, E. J. Douzery, A glimpse on the pattern of rodent diversification: a phylogenetic approach. BMC Evol. Biol. 14, 12–88 (2012).

41. 41. M. Halgren et al., The timescale and magnitude of 1/f aperiodic activity decrease with cortical depth in humans, macaques, and mice. bioRxiv [Preprint] (2021). https://www.biorxiv.org/content/10.1101/2021.07.28.454235v2 (accessed 26 July 2024).

42. R. D. Burwell, The Parahippocampal Region: Corticocortical Connectivity. Ann. N. Y. Acad. Sci. 911, 25–42 (2006).

43. P. Lavenex, D. G. Amaral, Hippocampal-neocortical interaction: a hierarchy of associativity. Hippocampus 10, 420–430 (2000).

44. G. M. G. Shepherd, The Synaptic Organization of the Brain (Oxford University Press, ed. 5, 2003).

45. M. M. Halassa, S. M. Sherman, Thalamocortical Circuit Motifs: A General Framework. Neuron 103, 762–770 (2019).

46. G. M. G. Shepherd, N. Yamawaki, Untangling the cortico-thalamo-cortical loop: cellular pieces of a knotty circuit puzzle. Nat. Rev. Neurosci. 22, 389–406 (2021).

47. R. Chaudhuri et al., A Large-Scale Circuit Mechanism for Hierarchical Dynamical Processing in the Primate Cortex. Neuron 88, 419–431 (2015).

48. C. C. Hilgetag, A. Goulas, ’Hierarchy’ in the organization of brain networks. Philos. Trans. R. Soc. Lond. B Biol. Sci. 375, 20190319 (2020).

49. V. J. Sydnor et al., Neurodevelopment of the association cortices: Patterns, mechanisms, and implications for psychopathology. Neuron 109, 2820–2846 (2021).

50. M. Demirtaş et al., Hierarchical Heterogeneity across Human Cortex Shapes Large-Scale Neural Dynamics. Neuron 101, 1181–1194.e13 (2019).

51. S. Li, X. J. Wang, Hierarchical timescales in the neocortex: Mathematical mechanism and biological insights. Proc. Natl. Acad. Sci. U.S.A. 119, e2110274119 (2022).

52. S. M. Bailes, D. E. P. Gomez, B. Setzer, L. D. Lewis, Resting-state fMRI signals contain spectral signatures of local hemodynamic response timing. Elife 12, e86453 (2023).

53. A. T. Campo et al., Thalamocortical interactions shape hierarchical neural variability during stimulus perception. iScience 27, 7 (2024).

54. K. Reinhold, A. D. Lien, M. Scanziani, Distinct recurrent versus afferent dynamics in cortical visual processing. Nat. Neurosci. 18, 1789–1797 (2015).

55. Z. V. Guo et al., Maintenance of persistent activity in a frontal thalamocortical loop. Nature 545, 181–186 (2017).

56. H. Zhou, R. J. Schafer, R. Desimone, Pulvinar-Cortex Interactions in Vision and Attention. Neuron 89, 209–220 (2016).

57. L. I. Schmitt et al., Thalamic amplification of cortical connectivity sustains attentional control. Nature 545, 219–223 (2017).

58. A. Hummos et al., Thalamic regulation of frontal interactions in human cognitive flexibility. PLoS Comput. Biol. 18, e1010500 (2022).

59. D. Joel, I. Weiner, The connections of the dopaminergic system with the striatum in rats and primates: an analysis with respect to the functional and compartmental organization of the striatum. Neuroscience 96, 451–474 (2000).

60. B. J. Hunnicutt et al., A comprehensive excitatory input map of the striatum reveals novel functional organization. Elife 5, e19103 (2016).

61. X. J. Wang, Decision Making in Recurrent Neuronal Circuits. Neuron 60, 215–234 (2008).

62. J. D. Murray, J. Jaramillo, X. J. Wang, Working Memory and Decision-Making in a Frontoparietal Circuit Model. J. Neurosci. 37, 12167–12186 (2017).

63. M. Stern, N. Istrate, L. Mazzucato, A reservoir of timescales emerges in recurrent circuits with heterogeneous neural assemblies. Elife 12, e86552 (2023).

64. J. J. Jun et al., Fully integrated silicon probes for high-density recording of neural activity. Nature 551, 232–236 (2017).

65. P. Shamash, M. Carandini, K. D. Harris, N.A. Steinmetz, A tool for analyzing electrode tracks from slice histology. bioRxiv [Preprint] (2018). https://www.biorxiv.org/content/10.1101/447995v1 (accessed 26 July 2024).

